# Ocular dominance-dependent binocular combination of monocular neuronal responses in macaque V1

**DOI:** 10.1101/2023.10.27.564359

**Authors:** Sheng-Hui Zhang, Xing-Nan Zhao, Shi-Ming Tang, Cong Yu

## Abstract

Primates rely on two eyes to see depth, while keeping a stable vision when one eye is closed. Although psychophysical and modeling studies have investigated how monocular signals are combined to form binocular vision, the corresponding neuronal mechanisms, especially in V1 where most neurons become binocular but with different eye preferences, are not well understood. Here we used two-photon calcium imaging to compare monocular and binocular responses of thousands of V1 superficial-layer neurons in three awake macaques. Under monocular stimulation, neurons preferring the stimulated eye responded substantially stronger than those preferring both eyes. However, under binocular stimulation, the responses of neurons preferring either eye were suppressed, and those preferring both eyes were enhanced, so that neuronal responses became similar regardless of eye preferences. A neuronally realistic model of binocular combination, which includes ocular dominance-dependent divisive interocular inhibition, and binocular summation, is proposed to account for these observations.

## Introduction

Human and non-human primates often use binocular disparity, or the differences between the retinal images of two eyes, to perceive depth (stereopsis). In the meantime, the brain maintains a stable perception of the visual world when one eye closes and the incoming light is halved. Much has been known regarding the neural foundations of stereopsis (Barlow, Blakemore, & Pettigrew, 1967; Henriksen, Tanabe, & Cumming, 2016; Parker, Smith, & Krug, 2016; Welchman, 2016; Read, 2021). Nevertheless, whether and how the neurons respond differently to monocular and binocular stimulations is less studied (Poggio & Fischer, 1977; Prince, Pointon, Cumming, & Parker, 2002; Dougherty, Cox, Westerberg, & Maier, 2019; Mitchell et al., 2022). Adding to the complexity is the fact that many V1 neurons, although responding to stimulations in both eyes, have various degrees of eye preferences (Hubel & Wiesel, 1962, 1968; Shatz & Stryker, 1978). Thus, a more complete picture of binocular combination of monocular responses would illustrate monocular and binocular responses of neurons at different eye preferences, which is the purpose of the current study.

Previous neurophysiological recording studies have revealed that overall the binocular responses of macaque V1 neurons are lower than the sum of monocular responses to each eye (Prince et al., 2002; Mitchell et al., 2022), but are stronger than monocular responses either when one eye was stimulated (Prince et al., 2002; Mitchell et al., 2022), or when neurons’ preferred eye was stimulated (Mitchell et al., 2022). Further, there is evidence that neurons’ eye preference may have important functional roles. For example, Dougherty et al. (2019) reported that the responses of monocular neurons are more likely suppressed than facilitated by binocular stimulation. As our results will suggest later, this conclusion is valid when the monocular baseline is either the sum of monocular responses, or the monocular responses of either eye. Dougherty et al. (2019) also reported similar responses of neurons with monocular and binocular preferences, as well as similar responses of binocular neurons to monocular and binocular stimulations, which are not supported by our data. In addition, Mitchell, Carlson, Westerberg, Cox, and Maier (2023) reported that binocular combination of monocular stimuli with different contrasts is also affected by neurons’ eye preference.

A big and more complete picture of binocular combination of monocular responses, as well as the impacts of eye preferences, would come from large samples of neurons in neighboring ocular dominance columns, so that the data can be quantitatively described with sufficient statistical power, and analyzed with computational modeling. Two-photon calcium imaging is capable of recording thousands of neurons simultaneously at the single-neuron resolution, which is especially suitable for this task. In this study, we used a two-photon calcium imaging setup, which had been adapted for recording with awake macaques (Li, Liu, Jiang, Lee, & Tang, 2017), to measure responses of V1 superficial-layer neurons to monocular and binocular stimulations. Furthermore, a neuronally realistic binocular combination model, which considers ocular dominance-dependent interocular divisive inhibition, as well as binocular summation, is proposed to explain the data.

## Results

We recorded responses of V1 superficial-layer neurons to monocular (contralateral and ipsilateral) and binocular stimulations in three awake, fixating macaques. The stimulus was a high-contrast (0.9) Gabor grating presented at various orientations and spatial frequencies (SFs). The recording was performed at two cortical depths of the same response field of view (FOV) in first two monkeys (MA & MB), and at one depth in the third monkey (MC) as the first two monkeys had displayed similar results at two depths (Fig. 1A). Imaging processing identified a total of 10,168 neurons. Among them, 9,390 (92.4%) were tuned to orientation, SF, or both. Results from these orientation- and/or SF-tuned neurons were used in the following data analysis.

**Figure 1.**
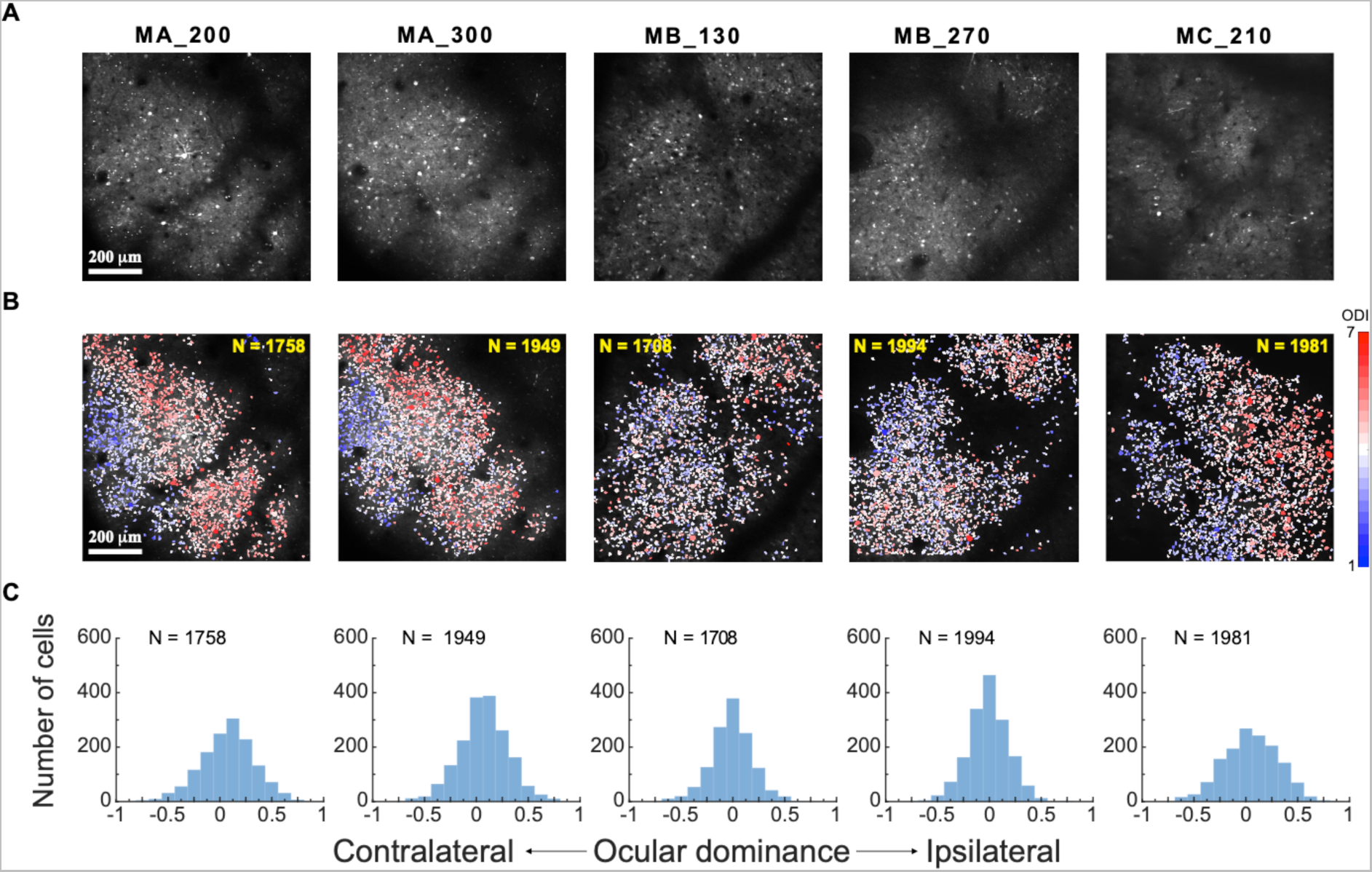
Eye preferences of V1 superficial-layer neurons in three macaques. **A**. Two-photon imaging. Average 2-photon images over a recording session for each response FOV. MA_200: Monkey A at a 200-μm cortical depth. **B**. Ocular dominance functional maps of each FOV/depth at single-neuron resolution. **C**. Frequency distributions of neurons of each FOV/depth as a function of ocular dominance index.

V1 superficial-layer neurons exhibited various degrees of eye preferences, consistent with the classical findings of Hubel and Wiesel (1962, 1968). An ocular dominance index (ODI) was calculated to characterize each neuron’s eye preference: ODI = (R_i_ – R_c_)/(R_i_ + R_c_), in which R_i_ and R_c_ were the neuron’s respective peak responses to ipsilateral and contralateral stimulations on the basis of data fitting (see Methods). Here ODI = −1 and 1 would indicate complete contralateral and ipsilateral eye preferences, respectively, and ODI = 0 would indicate equal preferences to both eyes. The ocular dominance functional maps at single-neuron resolution, especially those of Monkeys A and C, showed regions of neurons preferring either the contralateral (blue) or the ipsilateral eye (red), and transitional zones where neurons preferring both eyes (white) (Fig. 1B). The ocular dominance maps were similar at two cortical depths in Monkeys A and B, indicating possible ocular dominance columns (Fig. 1B). The frequency distributions of ODIs suggest more binocular neurons than monocular neurons in V1 superficial layers (Fig. 1C), similar to the normal distribution of ocularity index in Dougherty et al. (2019).

The responses of individual neurons were plotted against the ocular dominance index (ODI) when monocular stimulation was through the contralateral eye (Fig. 2A) or the ipsilateral eye (Fig. 2B). As expected, neurons with more negative ODIs responded stronger when the contralateral eye was stimulated, and those with more positive ODIs responded stronger when the ipsilateral eye was stimulated. The responses declined as neurons became more binocular. However, when both eyes were stimulated, the response differences among neurons of different eye preferences became not obvious (Fig. 2C).

**Figure 2.**
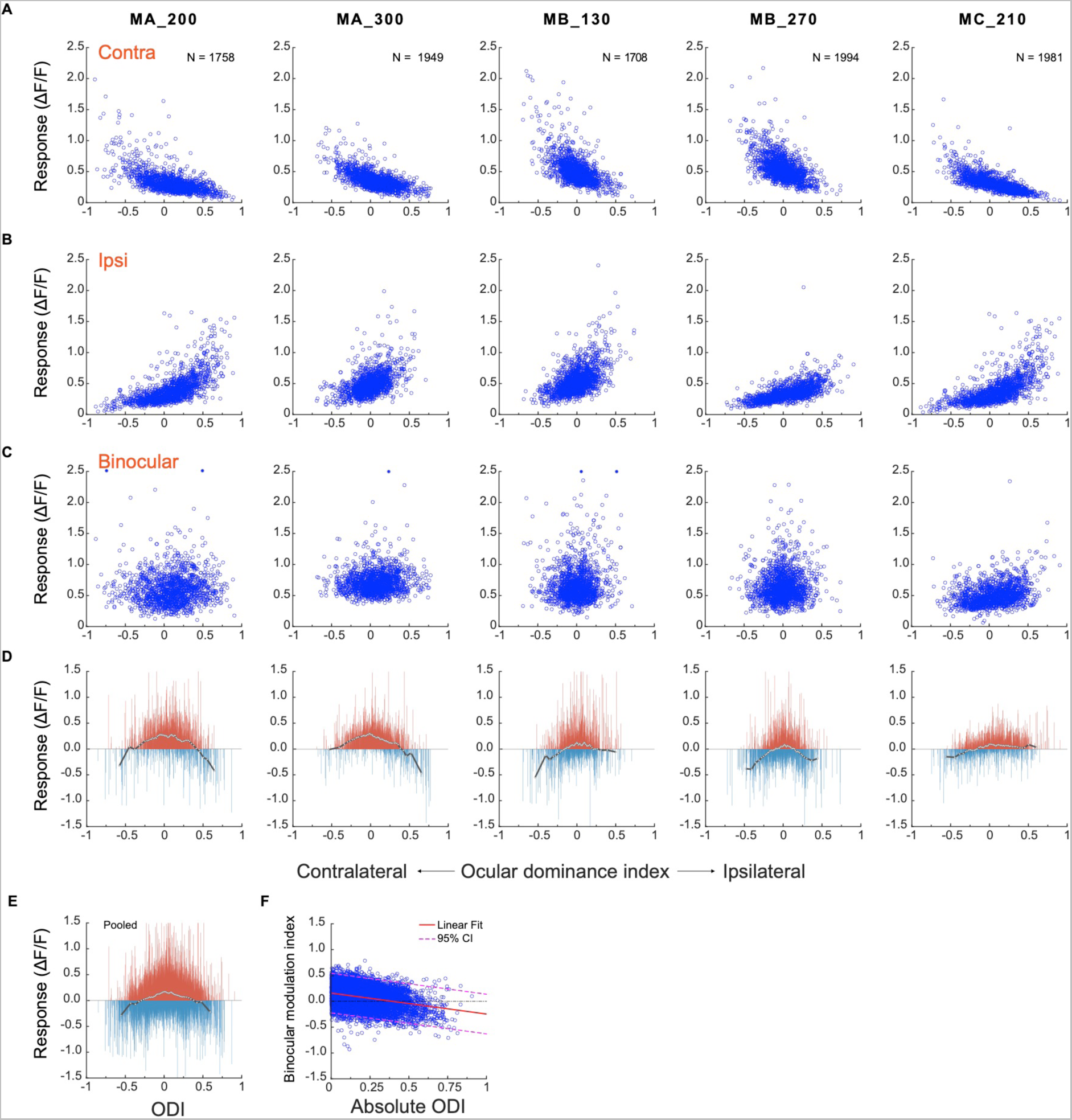
A comparison of neuronal responses to monocular and binocular stimulations. **A**. Responses of individual neurons against their ocular dominance indices with contralateral stimulation. **B**. Responses of the same neurons against their ocular dominance indices with ipsilateral stimulation. **C**. The binocular responses of the same neurons. **D**. The difference between binocular and monocular responses (R_b_ – max(R_i_, R_c_)). Each vertical line represents one neuron. To summarize the results, neurons of each FOV/depth are evenly divided into 60 bins in the order of the ocular dominance index. White dots represent the median responses of respective bins and are connected with a black line. **E**. The differences between binocular and monocular responses of individual neurons pooled over five FOVs/depths. **F**. Binocular modulation index as a function of absolute ODI and the linear fit. The binocular modulation index of each neuron was defined as (R_b_ – max (R_i_, R_c_))/(R_b_ + max (R_i_, R_c_)).

These trends can be better appreciated when the difference between binocular and monocular responses are presented for each FOV/depth in Fig. 2D: Under monocular stimulation, if only the neuronal responses to the preferred eye were taken into consideration (i.e., a neuron’s higher response to ipsilateral vs. contralateral stimulations), more monocular neurons (ODIs farther from 0) tended to have stronger responses than more binocular neurons (ODIs closer to 0). However, under binocular stimulation, the responses of more monocular neurons were suppressed, and those of more binocular neurons were enhanced, by binocular stimulation. These trends are best appreciated in pooled data over five FOVs/depths (Fig. 2E). Further, linear regression confirmed a significant dependence of binocular modulation on absolute ODI (y = −0.41x + 0.16, p <0.001) (Fig. 2E). That is, responses of neurons with lower absolute ODI (i.e., binocular neurons) tended to be more enhanced, and responses of neurons with higher absolute ODI (i.e., monocular neurons) tended to be more suppressed.

### Modeling monocular and binocular responses

We used the following steps to develop a model that can account for the current monocular responses and their binocular combination. Monocular and binocular data of each FOV/depth, as well as the pooled data, were first normalized by the respective median of the binocular responses of all neurons in the same FOV/depth. The normalized data were then divided into 60 bins in the order of the ocular dominance index, and the median values of 60 bins were used for model fitting (Figs. 3A-C).

**Figure 3.**
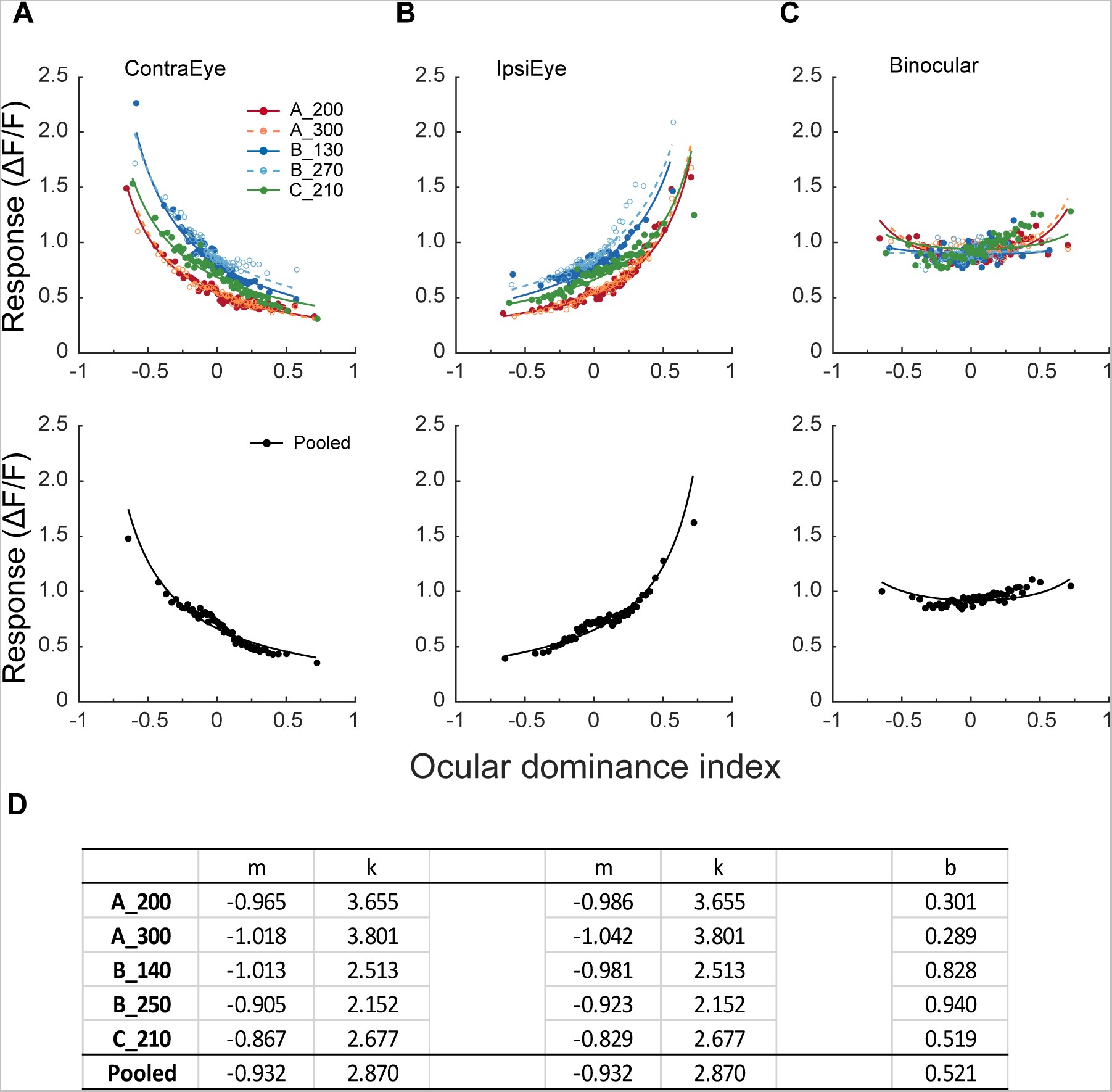
Modeling monocular and binocular responses. **A & B**. Median neuronal responses to contralateral and ipsilateral stimulations as a function of ocular dominance index and respective data fitting with Eq. 2. Neurons are evenly divided into 60 bins in the order of the ocular dominance index, with each bin containing 28-33 neurons that varied among different FOVs/depths (156 neurons for the pooled data). Each datum represents the median response of a bin. Free parameter k was kept equal during contralateral and ipsilateral data fitting. **C**. Binocular responses as a function of ocular dominance index and data fitting with Eq. 3 for the same bins of neurons. During binocular data fitting, parameters k, m_i_, and m_c_ were inherited from monocular data fitting, and only b was the free-changing parameter. **D**. The values of free parameters from monocular and binocular data fitting.

#### Monocular responses

First, a neuron’s monocular responses to contralateral and ipsilateral stimulations could be respectively described by a divisive gain control model:

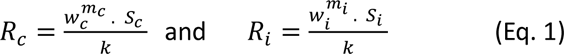

Here S_c_ and S_i_ were stimulus contrasts, w_c_ and w_i_ were linear transformations of a neuron’s ocular dominance index from [-1 1] to [0 1]: w_c_ = (ODI + 1)/2 and w_i_ = 1 – w_c_, m_c_ and m_i_ represented monocular nonlinearity, and k represented divisive gain control. Because S_c_ and S_i_ in our experiments were 0.9, which were about equal to 1 (full contrast) due to neuronal response saturation, the above equations were simplified as

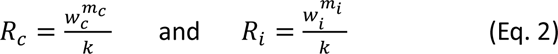

Eq. 2 was used to fit the binned median contralateral and ipsilateral data (Figs. 3A & B), with the parameter k being equal for contralateral and ipsilateral responses. The fitting revealed that m_c_ and m_i_ were negative and close to −1, which resulted in a quick decline of 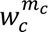 and a quick increase of 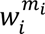 as a function of the ocular dominance index since w_c_ and w_i_ ∈ [0, 1]. The fit quality indices (Busse, Wade, & Carandini, 2009) ranged 0.92-0.94 for the contralateral condition (Fig. 3A) and 0.87-0.93 for the ipsilateral condition (Fig. 3B), suggesting adequate goodness of fit. The fitting parameters are listed in Fig. 3D.

#### Binocular responses

Fig. 2D-F earlier has indicated that neuronal responses to binocular stimulation change from suppression to enhancement as neurons’ ocular dominance changes from monocular to binocular, which may reflect the ocular dominance-dependent net effect of interocular suppression and binocular summation. Therefore, we added interocular response suppression to Eq. 2 by letting monocular responses from each eye be further normalized by an interocular suppression factor w_i_^b^ or w_c_^b^ (Eq. 3). In other words, the strength of interocular response suppression was decided by the linearly transformed ODI with a nonlinear exponent b. Finally, the normalized responses from two eyes were summed to simulate the binocular responses of neurons R_b_, completing the model of binocular combination (Eq. 3).

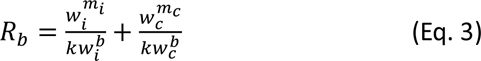

Although not shown in Eq. 3, we also assumed that the nonlinear exponent b also depends on the contrast of the stimulus presented to the other eye (i.e., S_c_ or S_i_). Consequently, when S_c_ or S_i_ = 0 under monocular stimulation, R_c_ or R_i_ = 0 (Eq. 1), and interocular suppression w_i_^b^ or w_c_^b^ = 1, so Eq. 3 changes back to Eq. 2. It is only when S_c_ and S_i_ are equal and close to 1, as in the current study, that interocular suppression and binocular combination would be in the current Eq. 3 format.

When fitting binocular responses, the parameters m_i_, m_c_, and k were inherited from earlier monocular data fitting and remained fixed. Only b was allowed to change. Data fitting resulted in flat binocular response functions (Fig. 3C) with satisfactory goodness-of-fit (fit quality index = 0.88 to 0.92).

The effects of interocular suppression and binocular summation in the model, as well as their contributions to the binocular response, may be better appreciated in Fig. 4. Fig. 4A uses Eqs. 2 and 3 and the parameters from fitting of pooled data (Fig. 3D) to simulate the contralateral (blue curve with label R_c_), ipsilateral (red curve with label R_i_), and binocular (black curve) response functions against the ocular dominance index. The arithmetic sum of contralateral and ipsilateral response functions was also simulated (grey dashed curve labeled R_i_ + R_c_). In addition, neuronal responses to preferred eye stimulation would consist of the higher branches of contralateral and ipsilateral response functions. It is apparent that binocular responses cannot be explained by the sum of monocular responses, as binocular responses are substantially lower than the summed monocular responses for both monocular and binocular neurons. Nor can binocular responses be explained by the responses to the preferred eye, as binocular responses are also lower than those to the preferred eye (the larger of the two monocular responses) for monocular neurons. Instead, the median of the binocular response function (black arrow) in each data set is close to but still more or less higher than the median of the contralateral (blue arrow) or ipsilateral response function (red arrow), which is consistent with previous reports (Prince et al., 2002; Mitchell et al., 2022).

**Figure 4.**
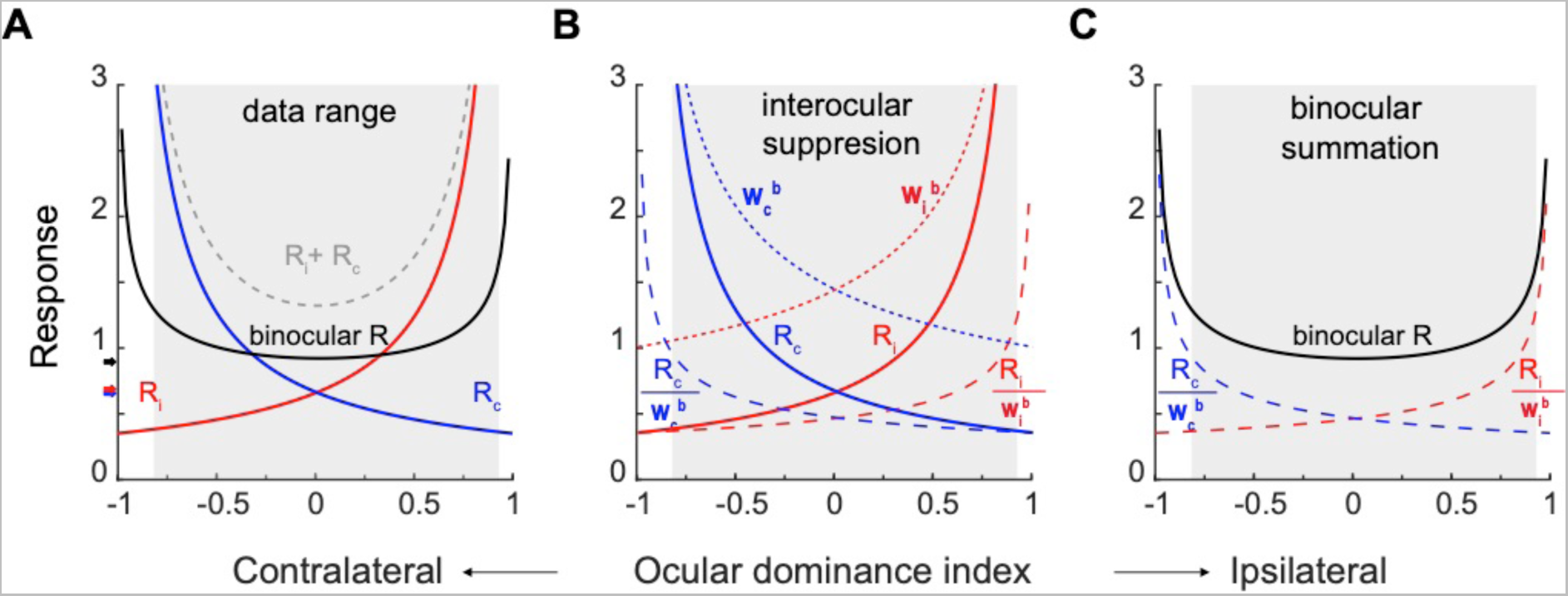
Monocular and binocular responses modeled with binocular suppression and binocular summation. **A.** The solid blue, red, and black curves are simulations of the contralateral, ipsilateral, and binocular responses on the basis of fitting of pooled data (Fig. 3D). The grey dashed curve simulates binocular responses as the arithmetic sum of contralateral and ipsilateral responses. The higher branches of contralateral and ipsilateral response functions represent monocular responses with preferred eye stimulation. The black, blue, and red arrows indicate the median binocular, contralateral, and ipsilateral responses, respectively, from pooled data. The shadowed area indicates the region where actual neurons existed on the basis of the ocular dominance index. **B.** Interocular suppression. The contralateral and ipsilateral responses (R_c_ & R_i_) are divided by respective interocular suppression factors W_c_^b^ and W^b^to produce interocular-suppressed responses R_c_/w_c_^b^ and R_i_/w_i_^b^. **C.** Binocular summation. R_c_/w_c_^b^ and R_i_/w_i_^b^ are summed to produce the final binocular responses R_b_.

Fig. 4B plots the interocular suppression factor w_c_^b^ (dotted blue curve) for the contralateral response R_c_ (solid blue curve from 4A), and w_i_^b^ (dotted red curve) for the ipsilateral response R_i_ (solid red curve from 4A). The interocular suppression factors w_c_^b^ and w_i_^b^ are larger with neurons that are more monocular than with neurons that are more binocular. The R_c_ and R_i_ are divided by the respective interocular suppression factor, producing the normalized contralateral responses (dashed blue curve labeled R_c_/w_c_^b^) and ipsilateral responses (dashed red curve labeled R_i_/w_i_^b^). These normalized curves are lower than original monocular responses, especially for neurons that are more monocular (ODIs farther from 0), showing interocular suppression.

Then the normalized monocular curves are summed up in Fig. 4C, which represents binocular summation and produces the final binocular responses (R_b_). Therefore, as a result of combined interocular suppression and binocular summation, the final curve for binocular responses becomes nearly flat within the data range.

## Discussion

The current study compared the responses of large samples of V1 superficial-layer neurons to monocular and binocular stimulations in three macaques. The monocular response functions show steep changes as a function of the ocular dominance index, but binocular response functions are largely flat, regardless of neurons’ eye preferences. Modeling efforts indicated that when signals from two eyes are combined, interocular divisive suppression, which is more prominent with neurons preferring one eye, and binocular summation, which is more prominent with binocular neurons, together produce the nearly flat binocular response function within a broad range of ocular dominance indices. These findings implicate that at least for neurons in superficial layers of V1, significant ocular dominance may result from a release of interocular suppression during monocular stimulation, an unusual viewing condition as our vision is typically binocular, rather than a lack of binocular combination of inputs from upstream monocular neurons.

We introduced this paper by citing the stable vision with monocular or binocular viewing. Relevant to this issue, Fig. 4A shows that the median of binocular neuronal responses is only slightly higher than that of monocular responses of either eye (black vs. red/blue arrows), consistent with earlier reports (Prince et al., 2002; Mitchell et al., 2022). The overall similar monocular and binocular responses may account for a large proportion of the stable vision under both viewing conditions. What is intriguing is our finding that the very small response changes actually comprise of dramatic binocular suppression with monocular neurons and facilitation with binocular neurons. Furthermore, slightly higher binocular responses suggest that they might be further modulated by additional mechanisms to achieve stable vision.

In Dougherty et al. (2019), the responses of monocular and binocular neurons are similar under monocular stimulation, and only monocular neurons, but not binocular neurons, are significantly suppressed by binocular stimulation. In contrast, our results show that monocular neurons respond to stimulation in the preferred eye much more strongly than binocular neurons do to the same monocular stimulation (Fig. 2C). Moreover, although confirming the observation of interocular suppression of monocular neurons responding to the preferred eye in Dougherty et al. (2019), our results also indicate enhanced responses of binocular neurons to binocular stimulation. Considering the large diversity of binocular responses of neurons with similar eye preferences (Fig. 2C), it is very possible to obtain inaccurate statistical estimates with small, and possibly biased, samples of neurons in electrode recording experiments. In addition, it is unclear whether the discrepancies are caused by different temporal resolutions of electrode recording and calcium imaging. The results of Dougherty et al. (2019) represent changes of neuronal spike activities over a period of approximately 50-200 ms after the stimulus onset, which may reflect the sustained neuronal responses to the stimulus and possible feedback signals. Calcium signals are much slower and indicative of the aggregated neuronal responses over a longer period (up to 1000 ms in the current study. They should have smeared, rather than exaggerated, the differences between monocular and binocular responses, though we cannot exclude the possibility that some neuronal response changes beyond 200 ms are responsible for the discrepancies.

Binocular combination of monocular signals has long been recognized to involve both interocular suppression and binocular summation (DeSilva & Bartley, 1930; Cohn & Lasley, 1976; Li & Atick, 1994). Most more recent models use divisive normalization to explain interocular suppression (e.g., Cogan, 1987; Anderson & Movshon, 1989; Ding & Sperling, 2006; Moradi & Heeger, 2009; Huang, Zhou, Zhou, & Lu, 2010; Ding, Klein, & Levi, 2013; Mitchell et al., 2023). Our modeling effort shows that a similar divisive interocular suppression and binocular summation model can account for neuronal response changes under monocular and binocular stimulations, with the distinction that the divisive interocular suppression is additionally controlled by a neuron’s ocular dominance. The latter also controls the responses of individual neurons to monocular stimulation, as shown in Figs. 3A-B. The critical roles of ocular dominance have been largely overlooked by extant binocular vision models to our knowledge, except that Anderson and Movshon (1989) demonstrated that a model consisting of multiple ocular dominance channels can better explain their psychophysical adaptation data, and that Mitchell et al. (2023) revealed that binocular combination of different contrasts presented to different eyes are affected by neurons’ ocularity preference. We hope that our two-photon imaging results can be incorporated into future neuronally plausible models of binocular vision.

On the basis of current findings, future two-photon imaging work shall compare neural responses to monocular and binocular stimulations with uneven effective stimulus contrasts due to physical contrast differences (Anderson & Movshon, 1989; Mitchell et al., 2023), monocular adaptation (Anderson & Movshon, 1989), and short-term monocular deprivation (Lunghi, Burr, & Morrone, 2011), as well as the relevant roles of ocular dominance of individual neurons. These results would help understand abnormal binocular vision in patients with strabismus and amblyopia. In addition, in our experiments, binocular stimuli were presented with zero disparity, which best triggered the responses of neurons with zero-disparity tuning. A more realistic model of binocular combination also requires the consideration of neurons with other disparity-tuning profiles.

### Limitations of the current study

Although capable of sampling a large number of neurons at cellular resolution and with low sampling bias, two-photon calcium imaging has its known limitations that may better make it a complementary research tool to electrophysiological recordings. For example, two-photon imaging can only sample neurons from superficial-layers, while binocular neurons also exist in deeper layers, and even neurons in the input layer are affected by feedback from downstream binocular neurons to exhibit binocular response properties (Dougherty et al., 2019). Furthermore, calcium signals are relatively slow and cannot reveal the fast dynamics of neuronal responses. Due to these spatial and temporal limitations, a more complete picture of the neuronal mechanisms underlying binocular combination of monocular responses may come from studies using both technologies.

In addition, calcium signals may exaggerate the nonlinear properties of neurons. Although calcium signals indicated by GCaMP5, our favored choice of calcium indicator, displays a linear relationship to neuronal spike rates within a range of 10-150 Hz (Li et al., 2017), weak and strong signals out of this range are more nonlinear, and may appear poorer and stronger, respectively, than electrode-recorded effects. Consequently, the differences in population responses between monocular and binocular stimulations revealed by this study might be less pronounced.

## Materials and Methods

### Monkey preparation

Monkey preparations were identical to those reported in a previous study (Guan, Ju, Tao, Tang, & Yu, 2021; Ju, Guan, Tao, Tang, & Yu, 2021). Three rhesus monkeys (*Macaca mulatta*) aged 4 to 6 years, respectively, were each prepared with two sequential surgeries under general anesthesia and strictly sterile condition. In the first surgery, a 20-mm diameter craniotomy was performed on the skull over V1. The dura was opened and multiple tracks of 100-150 nL AAV1.hSynap.GCaMP5G.WPRE.SV40 (AV-1-PV2478, titer 2.37e13 (GC/ml), Penn Vector Core) were pressure-injected at a depth of ∼350 μm. Then the dura was sutured, the skull cap was re-attached with three titanium lugs and six screws, and the scalp was sewn up. After the surgery, the animal was returned to the cage, treated with injectable antibiotics (Ceftriaxone sodium, Youcare Pharmaceutical Group, China) for one week. Postop analgesia was also administered. The second surgery was performed 45 days later. A T-shaped steel frame was installed for head stabilization, and an optical window was inserted onto the cortical surface. Data collection could start as early as one week later. More details of the preparation and surgical procedures can be found in Li et al. (2017). The procedures were approved by the Institutional Animal Care and Use Committee, Peking University.

### Behavioral task

After a ten-day recovery from the second surgery, monkeys were seated in primate chairs with head restraint. They were trained to hold fixation on a small white spot (0.1°) with eye positions monitored by an ISCAN ETL-200 infrared eye-tracking system (ISCAN Inc.) at a 120-Hz sampling rate (Monkeys A & B) or an Eyelink-1000 (SR Research) at a 1000-Hz sampling rate (Monkey C). During the experiment, trials with the eye position deviated 1.5° or more from the fixation before stimulus offset were discarded as ones with saccades and repeated. For the remaining trials, the eye positions were mostly concentrated around the fixation point, with eye positions in over 95% of trials within 0.5° from the fixation point. The viewing was binocular.

### Visual stimuli

For Monkeys A and B, visual stimuli were generated by the ViSaGe system (Cambridge Research Systems) and presented on a 21’’ Sony G520 CRT monitor (refresh rate = 80 Hz, resolution = 1280 pixel × 960 pixel, pixel size = 0.31 mm × 0.31 mm). Because of the space limit, the viewing distance and the monitor position varied depending on the stimulus spatial frequency (30 cm at 0.25, 0.5, and 1 cpd, 60 cm at 2 cpd, and 120 cm at 4 and 8 cpd). For Monkey C, visual stimuli were generated by Psychotoolbox 3 (Pelli & Zhang, 1991) and presented on a 27’’ Acer XB271HU LCD monitor (refresh rate = 80 Hz native, resolution = 2560 pixel × 1440 pixel native, pixel size = 0.23 mm × 0.23 mm). The viewing distance was 50 cm for lower frequencies (0.25 - 1 cpd) and 100 cm for higher frequencies (2 - 8 cpd). For both monitors, the screen luminance was linearized by an 8-bit look-up table, and the mean luminance was ∼47 cd/m^2^.

A drifting square-wave grating (SF = 4 cpd, contrast = full, speed = 3 cycles/s, starting phase = 0°, and size = 0.4° in diameter) was first used to determine the location, eccentricity (3.4° for Monkey A, 1.7° for Monkey B, and 1.1° for Monkey C) and size (0.8 - 1°) of the population receptive field associated with a recording field of view (FOV), as well as ocular dominance columns when monocularly presented to confirm the V1 location. This fast process used a 4 × objective lens mounted on the two-photon microscope and revealed no cell-specific information.

Neuronal responses were then measured with a high-contrast (0.9) Gabor grating (Gaussian-windowed sinusoidal grating) drifting at 2 cycles/s in opposite directions perpendicular to the Gabor orientation. The starting phase of the drafting Gabors was always 0°. The Gabor grating varied at 12 orientations from 0° to 165° in 15° steps, and 6 spatial frequencies from 0.25 to 8 cpd in 1-octave steps. In addition, three stimulus sizes (with constant stimulus centers) were used at each spatial frequency, for two purposes. First, our pilot measurements suggested very strong surround suppression with larger stimuli. Therefore, comparing the responses to different stimulus sizes helped approximate the RF size of each neuron that produced maximal response and least surround suppression. Second, for neurons whose RF centers and the stimulus center were misaligned, the RFs of some misaligned neurons might be better covered by larger stimuli. Still the RFs of additional neurons might have less overlap with the stimuli. These neurons would have weaker and less orientation-tuned responses because of the Gaussian-blurred stimulus edge, and would most likely be filtered out during our multiple steps of selection of orientation tuned neurons (see below). Specifically, the 0 of the Gaussian envelope of the Gabor were 0.64λ and 0.85λ at all spatial frequencies, and was additionally smaller at 0.42λ when spatial frequencies were 0.25 - 1 cpd, and larger at 1.06λ when spatial frequencies were 2 - 8 cpd (λ: wavelength; Gabors with the same σ in wavelength unit had the same number of cycles). Here at the smallest σ (0.42λ), the Gabors still had sufficient number of cycles (frequency bandwidths = 1 octave) (Graham, 1989), so that the actual stimulus spatial frequencies were precise at nominal values. In terms of visual angle, σ = 1.68°, 2.56°, and 3.36° at 0.25 cpd; 0.84°, 1.28°, and 1.68° at 0.5 cpd; 0.42°, 0.64°, and 0.85° at 1 cpd; 0.34°, 0.42°, and 0.53° at 2 cpd; 0.17°, 0.21°, and 0.26° at 4 cpd, and 0.08°, 0.11°, and 0.13° at 8 cpd, respectively.

Each stimulus was presented for 1000 ms, with an inter-stimulus interval (ISI) of 1500 ms that is sufficient to allow the calcium signals back to the baseline level (Guan, Zhang, Zhang, Tang, & Yu, 2020). Each stimulus condition was repeated 12 times with six repeats for each opposite direction. When a stimulus was presented monocularly to one eye, the other eye was covered with a translucent eye patch to reduce the impacts of short-term monocular deprivation. For Monkey A, binocular recording preceded monocular recording in separate days. During monocular recording contralateral and ipsilateral stimulations alternated in blocks of trials with at least 10-minute breaks, during which the eye mask was taken off. Recording at a specific viewing distance was completed with all trials at relevant SFs pseudo-randomly presented before proceeding to the next distance. For Monkeys B and C, binocular and monocular recordings were mixed and completed in two daily sessions. At a specific viewing distance, all binocular trials at relevant SFs were completed first, then contralateral and ipsilateral trials were completed in alternating blocks of trials with at least 10-minute eye-patch-off breaks. Again, recordings at a specific viewing distance were completed before proceeding to a different distance. Each block of trials typically lasted 20-25 minutes, but for Monkeys A & B some blocks involving three SFs could last up to 45 minutes. The strength of fluorescent signals (mean luminance of a small area) was monitored and adjusted if necessary for drift of fluorescent signals. We compared the response ratios of the last 2 trials over the first 2 trials for each stimulus condition in these extended blocks with ipsilateral and contralateral stimulations. The respective mean ratios were 0.94 and 0.86, suggesting that the recorded neuronal responses remained largely stable over extended blocks of trials.

### Two-photon calcium imaging

Two-photon calcium imaging was performed with a Prairie Ultima IV (In Vivo) two-photon microscope (Prairie Technologies) on Monkeys A and B, or a FENTOSmart two-photon microscope (Femtonics) on Monkey C, and a Ti:sapphire laser (Mai Tai eHP, Spectra Physics). GCaMP5 was chosen as the indicator of calcium signals because the fluorescence activities it expresses are linearly proportional to neuronal spike activities within a wide range of firing rates from 10-150 Hz (Li et al., 2017). One FOV of 850 x 850 μm^2^ was selected in each animal and imaged using a 1000-nm femtosecond laser under a 16 × objective lens (0.8 N.A., Nikon) at a resolution of 1.6 μm/pixel. Fast resonant scanning mode (32 frames per second) was chosen to obtain continuous images of neuronal activity (8 frames per second after averaging every 4 frames). Recording was first performed at a shallower depth, and some neurons with high brightness or unique dendrite patterns were selected as landmarks. In the next daily session, the same FOV at the same depth was located before recording with the help of landmarks, and the depth plane was lowered if recording was performed at a deeper depth (Monkeys A & B). Because of the time limit, recordings at a specific FOV/depth with monocular and binocular stimulations were completed in 2-3 consecutive daily sessions, but the same neurons could be precisely tracked over multiple recording sessions with the use of multiple landmark cues.

### Imaging data analysis: Initial screening of ROIs

Data were analyzed with customized MATLAB codes. A normalized cross-correlation based translation algorithm was used to reduce motion artifacts (Li et al., 2017). Then fluorescence changes were associated with corresponding visual stimuli through the time sequence information recorded by Neural Signal Processor (Cerebus system, Blackrock Microsystem). By subtracting the mean of the 4 frames before stimuli onset (*F0*) from the average of the 6th-9th frames after stimuli onset (*F*) across 5 or 6 repeated trials for the same stimulus condition (same orientation, spatial frequency, size, and drifting direction), the differential image (*ΔF = F -F0*) was obtained.

For a specific FOV at a specific recording depth, the regions of interest (ROIs) or possible cell bodies were decided through sequential analysis of 216 differential images in the order of spatial frequency (6), size (3), and orientation (12) (6 x 3 x 12 = 216). The first differential image was filtered with a band-pass Gaussian filter (size = 2-10 pixels), and connected subsets of pixels (>25 pixels, which would exclude smaller vertical neuropils) with average pixel value > 3 standard deviations of the mean brightness were selected as ROIs.

Then the areas of these ROIs were set to mean brightness in the next differential image before the bandpass filtering and thresholding were performed (This measure gradually reduced the SDs of differential images and facilitated detection of neurons with relatively low fluorescence responses). If a new ROI and an existing ROI from the previous differential image overlapped, the new ROI would be on its own if the overlapping area OA < 1/4 ROI_new_, discarded if 1/4 ROI_new_ < OA < 3/4 ROI_new_, and merged with the existing ROI if OA > 3/4 ROI_new_. The merges would help smooth the contours of the final ROIs. This process went on through all 512 differential images twice to select ROIs. Finally, the roundness for each ROI was calculated as:

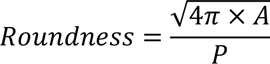

where 𝐴 was the ROI’s area, and 𝑃 was the perimeter. Only ROIs with roundness larger than 0.9, which would exclude horizontal neuropils, were selected for further analysis.

### Imaging data analysis: Orientation tuning, SF tuning, and ocular dominance

The ratio of fluorescence change (*ΔF/F0*) was calculated as a neuron’s response to a specific stimulus condition. For a specific cell’s response to a specific stimulus condition, the *F0_n_* of the n-th trial was the average of 4 frames before stimulus onset, and F_n_ was the average of 5th-8th frames after stimulus onset. F0_n_ was then averaged across 12 trials to obtain the baseline F0 for all 12 trials (for the purpose of reducing noises in the calculations of responses), and ΔF_n_/F0 = (F_n_-F0)/F0 was taken as the neuron’s response to this stimulus at this trial. The final response was averaged over 11 trials, minusing the 12^th^ trial that showed the weakest and often negative response. For a small portion of neurons (e.g., ∼3% in Monkeys A, ∼8% in monkey B and ∼2% in Monkey C) showing direction selectivity as their responses to two opposite directions differed significantly (p < 0.05, Friedman test), the 6 trials at the preferred direction was considered for calculations of ΔF_n_/F0 as the cell’s responses to a particular stimulus. F0 was still averaged over 12 trials at two opposite directions.

Several steps were then taken to decide whether a neuron was tuned to orientation and/or spatial frequency, and if so, its ocular dominance index. For each monocular condition, first the orientation, SF, and size (σ) producing the maximal response among all conditions were selected. Then responses to other 11 orientations and 5 SFs were decided at the selected SF and size. Second, to select orientation and/or SF tuned neurons, a non-parametric Friedman test was performed to test whether a neuron’s responses at 12 orientations or 6 SFs were significantly different from each other at least under one monocular stimulation. To reduce Type-I errors, the significance level was set at α = 0.01.

Third, for those showing significant orientation difference, the trial-based orientation responses of each neuron were fitted with a Gaussian model with a MATLAB nonlinear least-squares function: lsqnonlin:

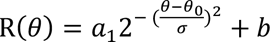

where R(θ) was the response at orientation θ, free parameters a_1_, θ_0_, σ, and b were the amplitude, peak orientation, standard deviation of the Gaussian function, and minimal response of the neuron, respectively. Only neurons with goodness of fit R_2_ > 0.5 at least under one stimulation condition were finally selected as orientation tuned neurons. Fourth, for those showing significant SF difference, the trial-based SF responses of each neuron were further fitted with a Difference-of-Gaussian model.

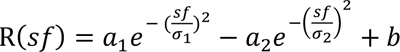

where R(sf) was a neuron’s response at spatial frequency sf, free parameters a_1_, σ_1_, a_2_, and σ_2_ were amplitudes and standard deviations of two Gaussians, respectively, and b was the minimal response among 6 spatial frequencies. Only those with goodness of fit R^2^ > 0.5 at least under one monocular stimulation were selected as SF tuned neurons.

The ocular dominance index (ODI) was calculated to characterize each orientation and/or SF tuned neuron’s eye preference: ODI = (R_i_ – R_c_)/(R_i_ + R_c_), in which R_i_ and R_c_ were the neuron’s respective peak responses at the best orientation and SF to ipsilateral and contralateral stimulations on the basis of data fitting. Here ODI = −1 and 1 would indicate complete contralateral and ipsilateral eye preferences, respectively, and ODI = 0 would indicate equal preferences to both eyes.

### Model fitting

Monocular and binocular data in Fig. 4 were fitted by Eqs. 2 and 3, respectively. The goodness-of-fit was indicated by a fit quality index q with a range of 0-1, which is the root mean square deviation between the observed responses and the model normalized by the observed response mean (Busse et al., 2009):

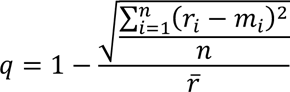

where *i* indicates the i_th_ bin, *r* indicates median response of a specific bin, and m indicates the corresponding model predictions.

## Acknowledgments

This study was supported by the National Science and Technology Innovation 2030 Major Program (2022ZD0204600), Natural Science Foundation of China grants 31230030 and 31730109, and funds from Peking-Tsinghua Center for Life Sciences, Peking University.

## Competing interests

None.

## Notes

### Competing Interest Statement

The authors have declared no competing interest.

### Summary of Updates

Revision is made on the basis of reviewers' comments.

## References

Anderson, P. A., & Movshon, J. A. (1989). Binocular combination of contrast signals. Vision Res, 29, 1115–1132.

Barlow, H. B., Blakemore, C., & Pettigrew, J. D. (1967). The neural mechanism of binocular depth discrimination. J Physiol, 193, 327–342.

Busse, L., Wade, A. R., & Carandini, M. (2009). Representation of concurrent stimuli by population activity in visual cortex. Neuron, 64, 931–942.

Cogan, A. I. (1987). Human binocular interaction: towards a neural model. Vision Res, 27, 2125–2139.

Cohn, T. E., & Lasley, D. J. (1976). Binocular vision: two possible central interactions between signals from two eyes. Science, 192, 561–563.

DeSilva, H. R., & Bartley, S. H. (1930). Summation and subtraction of brightness in binocular perception. British Journal of Psychology, 20, 242–252.

Ding, J., Klein, S. A., & Levi, D. M. (2013). Binocular combination of phase and contrast explained by a gain-control and gain-enhancement model. J Vis, 13, 13.

Ding, J., & Sperling, G. (2006). A gain-control theory of binocular combination. Proc Natl Acad Sci U S A, 103, 1141–1146.

Dougherty, K., Cox, M. A., Westerberg, J. A., & Maier, A. (2019). Binocular Modulation of Monocular V1 Neurons. Curr Biol, 29, 381–391 e384.

Graham, N. (1989). Visual Pattern Analyzers (Oxford Psychology Series, No. 16). New York: Oxford University Press.

Guan, S. C., Ju, N., Tao, L., Tang, S. M., & Yu, C. (2021). Functional organization of spatial frequency tuning in macaque V1 revealed with two-photon calcium imaging. Progress in Neurobiology, 205, 102120.

Guan, S. C., Zhang, S. H., Zhang, Y. C., Tang, S., & Yu, C. (2020). Plaid detectors in macaque V1 revealed by two-photon calcium imaging. Current Biology, 30, 934–940.

Henriksen, S., Tanabe, S., & Cumming, B. (2016). Disparity processing in primary visual cortex. Philos Trans R Soc Lond B Biol Sci, 371.

Huang, C. B., Zhou, J., Zhou, Y., & Lu, Z. L. (2010). Contrast and phase combination in binocular vision. PLoS One, 5, e15075.

Hubel, D. H., & Wiesel, T. N. (1962). Receptive fields, binocular interaction and functional architecture in the cat’s visual cortex. J Physiol, 160, 106–154.

Hubel, D. H., & Wiesel, T. N. (1968). Receptive fields and functional architecture of monkey striate cortex. J Physiol, 195, 215–243.

Ju, N., Guan, S. C., Tao, L., Tang, S. M., & Yu, C. (2021). Orientation tuning and end-stopping in macaque V1 studied with two-photon calcium imaging. Cerebral Cortex, 31, 2085– 2097.

Li, M., Liu, F., Jiang, H., Lee, T. S., & Tang, S. (2017). Long-term two-photon imaging in awake macaque monkey. Neuron, 93, 1049–1057 e1043.

Li, Z. P., & Atick, J. J. (1994). Efficient Stereo Coding in the Multiscale Representation. Network-Computation in Neural Systems, 5, 157–174.

Lunghi, C., Burr, D. C., & Morrone, C. (2011). Brief periods of monocular deprivation disrupt ocular balance in human adult visual cortex. Curr Biol, 21, R538–539.

Mitchell, B. A., Carlson, B. M., Westerberg, J. A., Cox, M. A., & Maier, A. (2023). A role for ocular dominance in binocular integration. Curr Biol, 33, 3884–3895 e3885.

Mitchell, B. A., Dougherty, K., Westerberg, J. A., Carlson, B. M., Daumail, L., Maier, A., & Cox, M. A. (2022). Stimulating both eyes with matching stimuli enhances V1 responses. iScience, 25, 104182.

Moradi, F., & Heeger, D. J. (2009). Inter-ocular contrast normalization in human visual cortex. J Vis, 9, 13 11–22.

Parker, A. J., Smith, J. E., & Krug, K. (2016). Neural architectures for stereo vision. Philos Trans R Soc Lond B Biol Sci, 371.

Pelli, D. G., & Zhang, L. (1991). Accurate control of contrast on microcomputer displays. Vision Res, 31, 1337–1350.

Poggio, G. F., & Fischer, B. (1977). Binocular interaction and depth sensitivity in striate and prestriate cortex of behaving rhesus monkey. J Neurophysiol, 40, 1392–1405.

Prince, S. J., Pointon, A. D., Cumming, B. G., & Parker, A. J. (2002). Quantitative analysis of the responses of V1 neurons to horizontal disparity in dynamic random-dot stereograms. J Neurophysiol, 87, 191–208.

Read, J. C. A. (2021). Binocular vision and stereopsis across the animal kingdom. Annu Rev Vis Sci, 7, 389–415.

Shatz, C. J., & Stryker, M. P. (1978). Ocular dominance in layer IV of the cat’s visual cortex and the effects of monocular deprivation. J Physiol, 281, 267–283.

Welchman, A. E. (2016). The Human Brain in Depth: How We See in 3D. Annu Rev Vis Sci, 2, 345–376.

